# May the phage be with you? Prophage-like elements in the genomes of Soft Rot *Pectobacteriaceae* (*Pectobacterium* spp. and *Dickeya* spp.)

**DOI:** 10.1101/399675

**Authors:** Robert Czajkowski

## Abstract

Soft Rot *Pectobacteriaceae* (SRP; *Pectobacterium* spp. and *Dickeya* spp., formerly known as pectinolytic *Erwinia* spp.) are necrotrophic bacterial pathogens infecting large number of plant species worldwide including agriculturally-important crops. Regardless of the SRP importance in agriculture, little is known about the bacteriophages infecting *Pectobacterium* spp. and *Dickeya* spp. and even less about prophages present in SRP genomes. Prophages are recognized as factors underlying bacterial virulence, genomic diversification and ecological fitness and have association with the novel phenotypic properties of bacterial hosts. Likewise, they are recognized as a driving force of bacterial evolution. The aim of this study was to analyze *Pectobacterium* spp. and *Dickeya* spp. complete genome sequences downloaded from GenBank (NCBI) for the presence of prophage sequences following their identification and (comparative) characterization with the main focus on current and upcoming perspectives in that field.

## INTRODUCTION

It is generally accepted that phages are the most abundant entities in the environment with an estimated number of 10^31^ particles on Earth. Consequently, they are present virtually in all surroundings where bacteria exists (Suttle, 2007). Based on their overall relationship with the host, they can be divided into virulent (lytic) and temperate (lysogenic) (Ackermann, 2003). Temperate bacteriophages integrates their genetic material into the host genome and persist inside bacterial cells as so-called prophages (Weinbauer, 2004). After integration prophages undergo the nonlytic growth mode named lysogenic state (Canchaya et al., 2004). During lysogeny phage DNA remains inactive excluding some regulatory and accessory genes, which are expressed in the host to maintain the dormant state of the virus. This dormant bacteriophage DNA may constitute even up to 20% of the host genomes (Casjens, 2003).

The present of prophages has a tremendous impact of bacterial fitness (Bondy-Denomy and Davidson, 2014;Nanda et al., 2014). They can induce host variability and evolution as well as prophages may influence adaptation of their hosts to the specific ecological niches (Wang et al., 2010;Fortier and Sekulovic, 2013;Varani et al., 2013). The presence of prophages may affect the bacterial genomes in several ways. For example, they are responsible for gene disruption or relocation which may lead to host phenotypic changes. Likewise, prophages may introduce new factors including pathogenicity determinants altering bacterium fitness and due to the switch between lytic and lysogenic cycle may further protect populations of the bacterial hosts in particular environment (e.g. in biofilms) (Brüssow et al., 2004). Consequently, prophages have been extensively studied in a number of bacterial species including plant pathogens to understand their role in bacterial ecology (Casjens, 2003;Varani et al., 2013).

Plant pathogenic Soft Rot *Pectobacteriaceae* (SRP) (Adeolu et al., 2016) (*Pectobacterium* spp. and *Dickeya* spp., formerly characterized as pectinolytic *Erwinia* spp. (Pérombelon, 2002)) are classified among the top ten the most important phytopathogens in agriculture (Mansfield et al., 2012). They cause significant losses in crop production (up 40% depending on the weather conditions, plant susceptibility and pathogen inoculum) i.e. in potato, carrot, tomato, onion, pineapple, maize, rice, hyacinth, chrysanthemum and calla lily worldwide (Perombelon and Kelman, 1980;Charkowski, 2018). SRP are widespread in various ecological niches *viz*. bulk and rhizosphere soils, water, sewage, on host and non-host plants as well as they are present on and inside insects (Perombelon and Kelman, 1980;Grenier et al., 2006;Rossmann et al., 2018). Likewise, SRP, because of their lifestyle, are often transferred between different environments (for example: from plant to soil, from plant to plant, from surface and/or irrigation water to plant, from water to soil, and vice verse) (Charkowski, 2018). In all these surroundings the bacteria can encounter lytic and temperate bacteriophages and hence may become easily repeatedly infected (Canchaya et al., 2004). It remains unknown however how latent infection with bacteriophages, resulting in viral genome integration to the host genome, influences the ecological fitness of *Pectobacterium* spp. and *Dickeya* spp. and bacterial adaptation and evolution.

The knowledge on prophages present in SRP genomes is very limited as till present only several temperate bacteriophages infecting *Pectobacterium* spp. and *Dickeya* spp. have been characterized (Varani et al., 2013;Czajkowski, 2015). These include temperate bacteriophage ϕEC2 infecting *D. dadantii* and *D. solani* (Resibois et al., 1984), bacteriophage ZF40 (Korol and Tovkach, 2012) infecting *P. carotovorum* subsp. *carotovorum*, bacteriophages phiTE (Blower et al., 2012), ECA29 and ECA41 (Evans et al., 2010) infecting *P. atrosepticum* and lytic/temperate bacteriophages LIMEstone 1 and LIMEstone2 infecting *D. solani* (Adriaenssens et al., 2012). Likewise, the biological role of only two prophages present in SRP genomes: ECA29 and ECA41 localized in the genome of *P. atrosepticum* strain SCRI1043 were fully elucidated so far (Evans et al., 2010).

The aim of this work was to assess the known, complete genome sequences of *Pectobacterium* spp. and *Dickeya* spp. strains present in GenBank (NCBI) for the occurrence of prophage-like sequences and their characterization with the genetic and (comparative) genomic tools. The implications of prophage presence in SRP genomes and the way how these genetic elements may contribute to ecological fitness of *Pectobacterium* spp. and *Dickeya* spp. are discussed.

## MATERIALS AND METHODS

### *Data collection and identification of candidate prophage sequences in complete genomes of* Dickeya *spp*. and Pectobacterium *spp.*

The identification and characterization strategy to find prophages in SRP genomes is presented in Figure 1. Fifty seven complete genome sequences (17 *Pectobacterium* spp. genomes and 40 *Dickeya* spp. genomes) were downloaded from NCBI (National Center for Biotechnology Information, https://www.ncbi.nlm.nih.gov/) (August 2018) (Table 1). Candidate prophage-like elements were identified using PHASTER (http://phaster.ca/) (Arndt et al., 2016) and PhiSpy (https://edwards.sdsu.edu/PhiSpy/index.php) (Akhter et al., 2012), following the manual inspection of the resulting sequences for the presence of attachment sites (*att*), gene(s) coding for integrase(s) and the genetic content of the prophage integration places as suggested by others (Boyd and Brüssow, 2002). Prophages were characterized on the basis of their homology with known phage sequences deposited in GenBank (https://www.ncbi.nlm.nih.gov/genbank/). The presence of structural genes in prophage sequences was verified by the VirFam (http://biodev.cea.fr/virfam/) (Lopes et al., 2014).

**Table 1.**
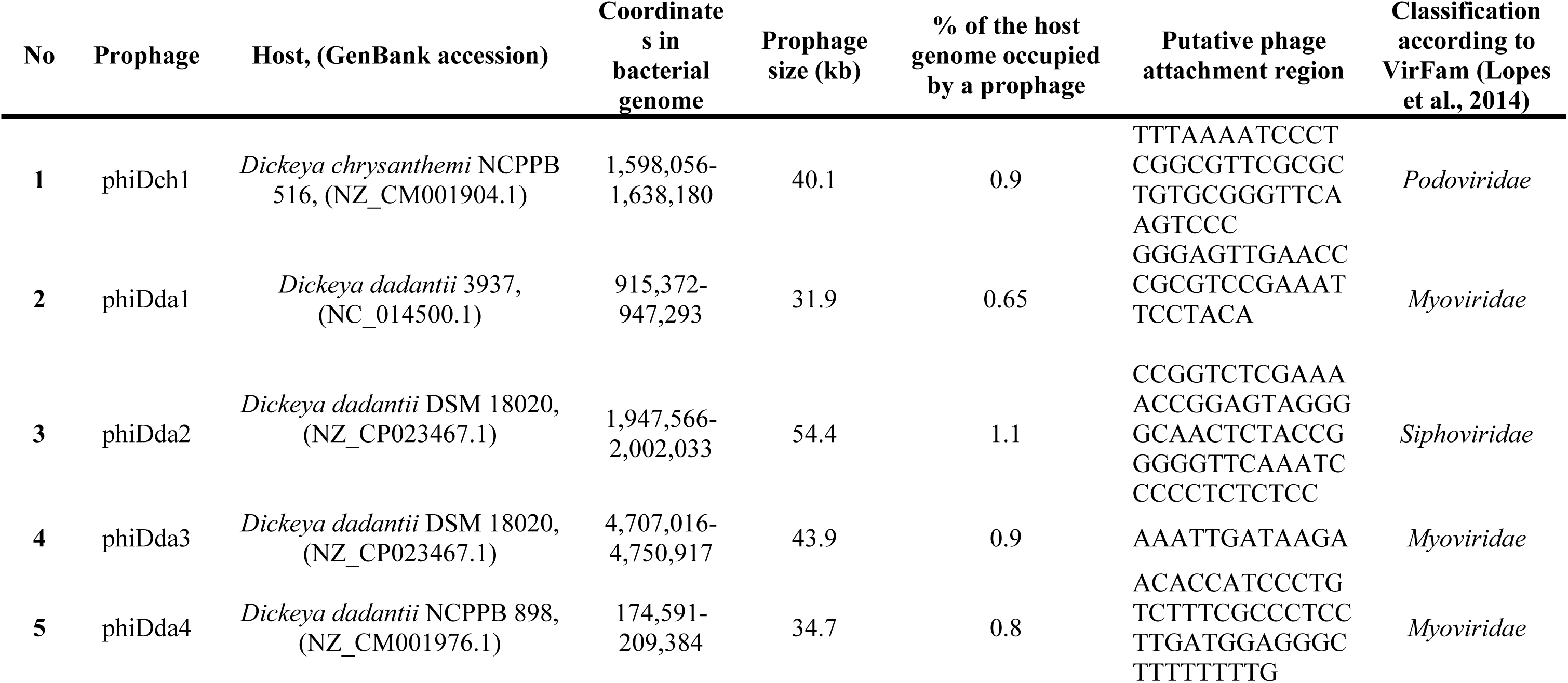
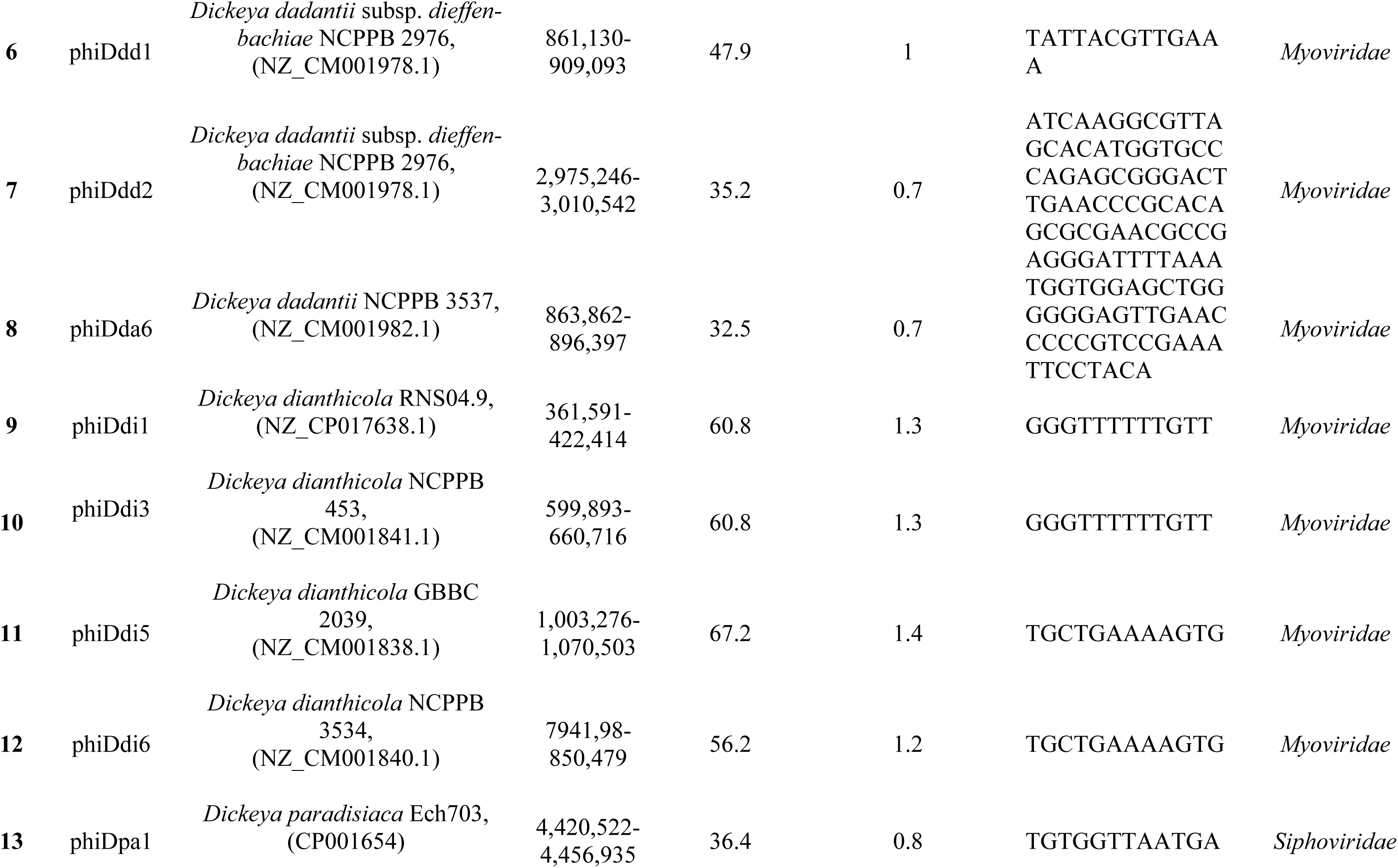
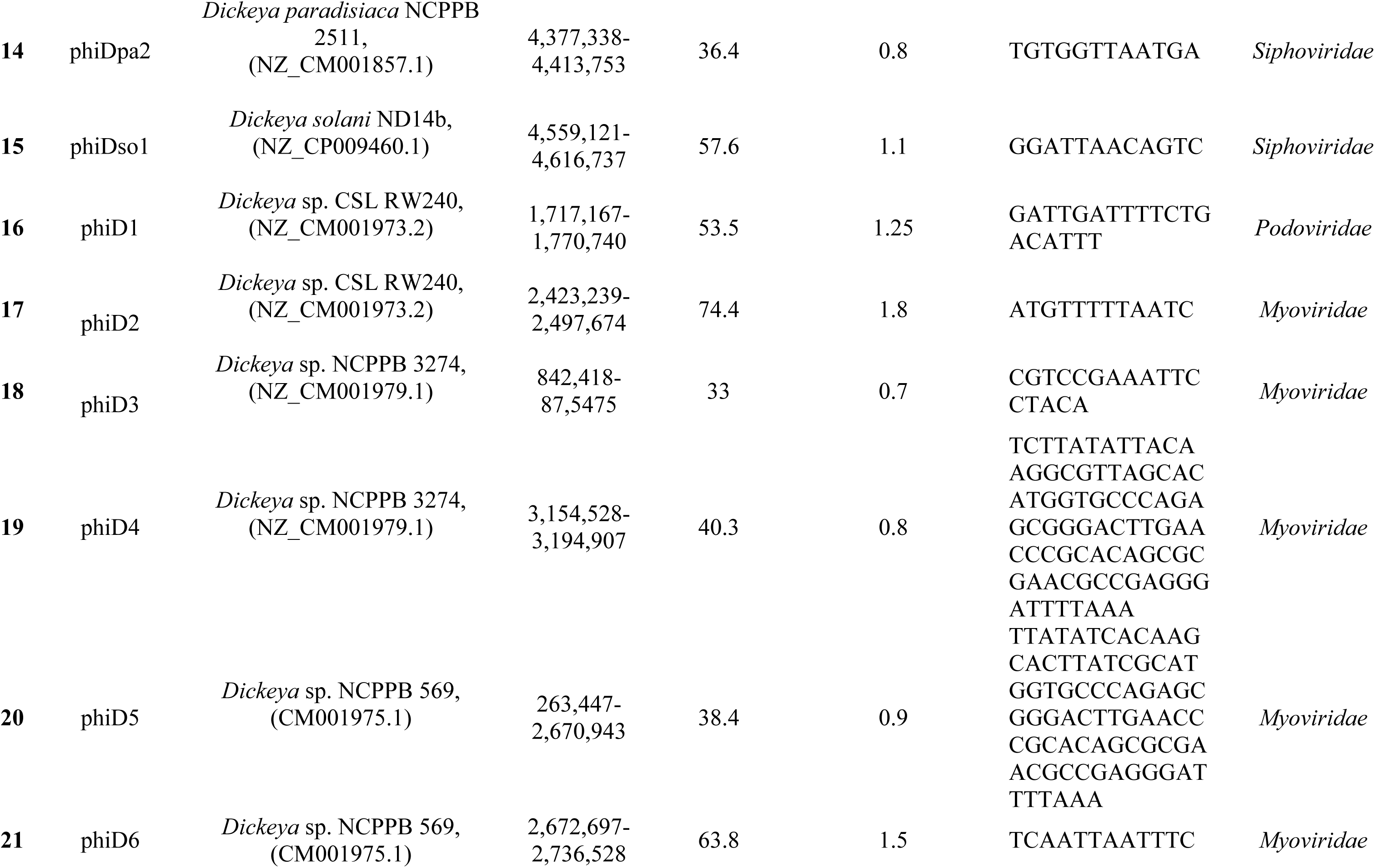
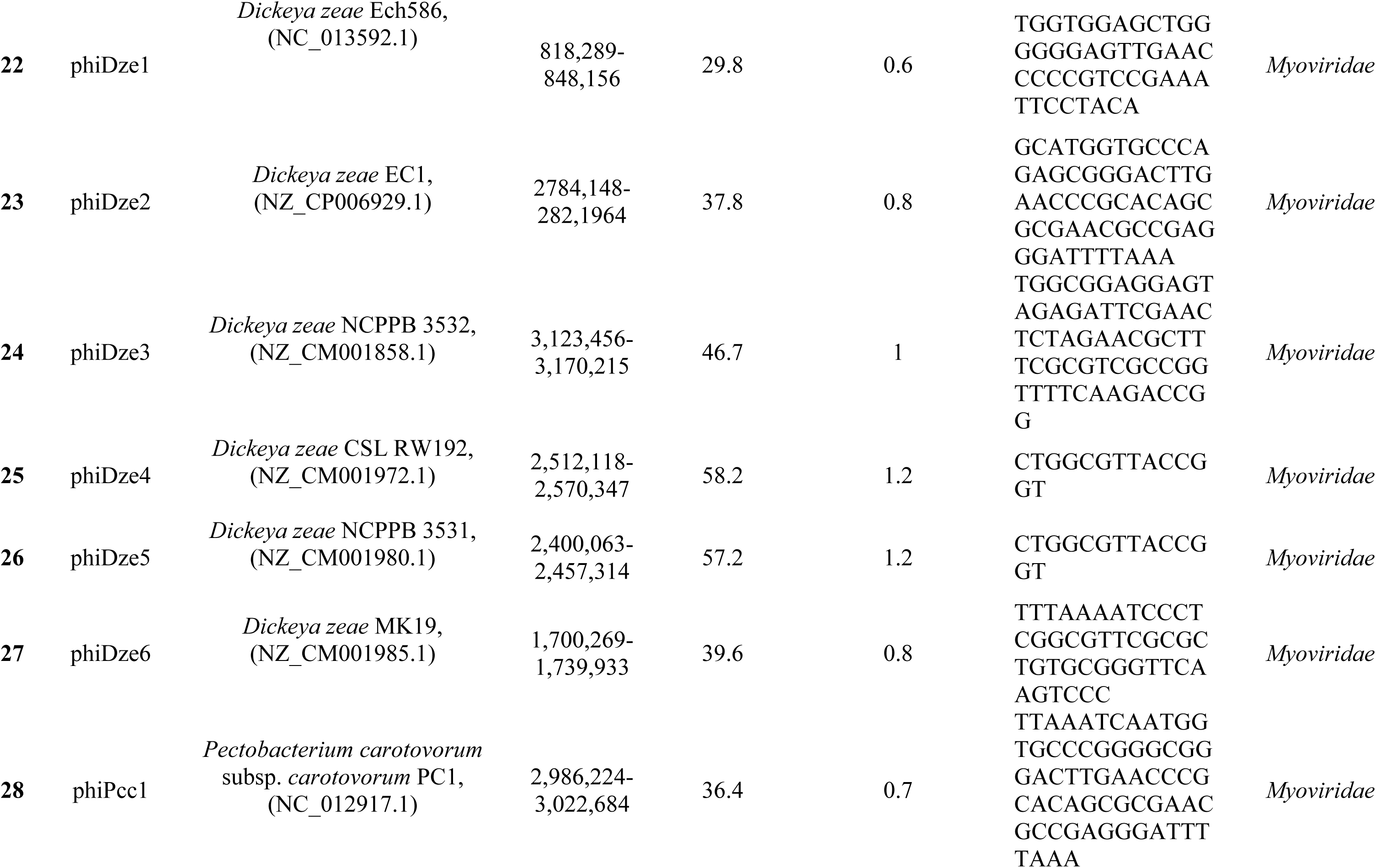
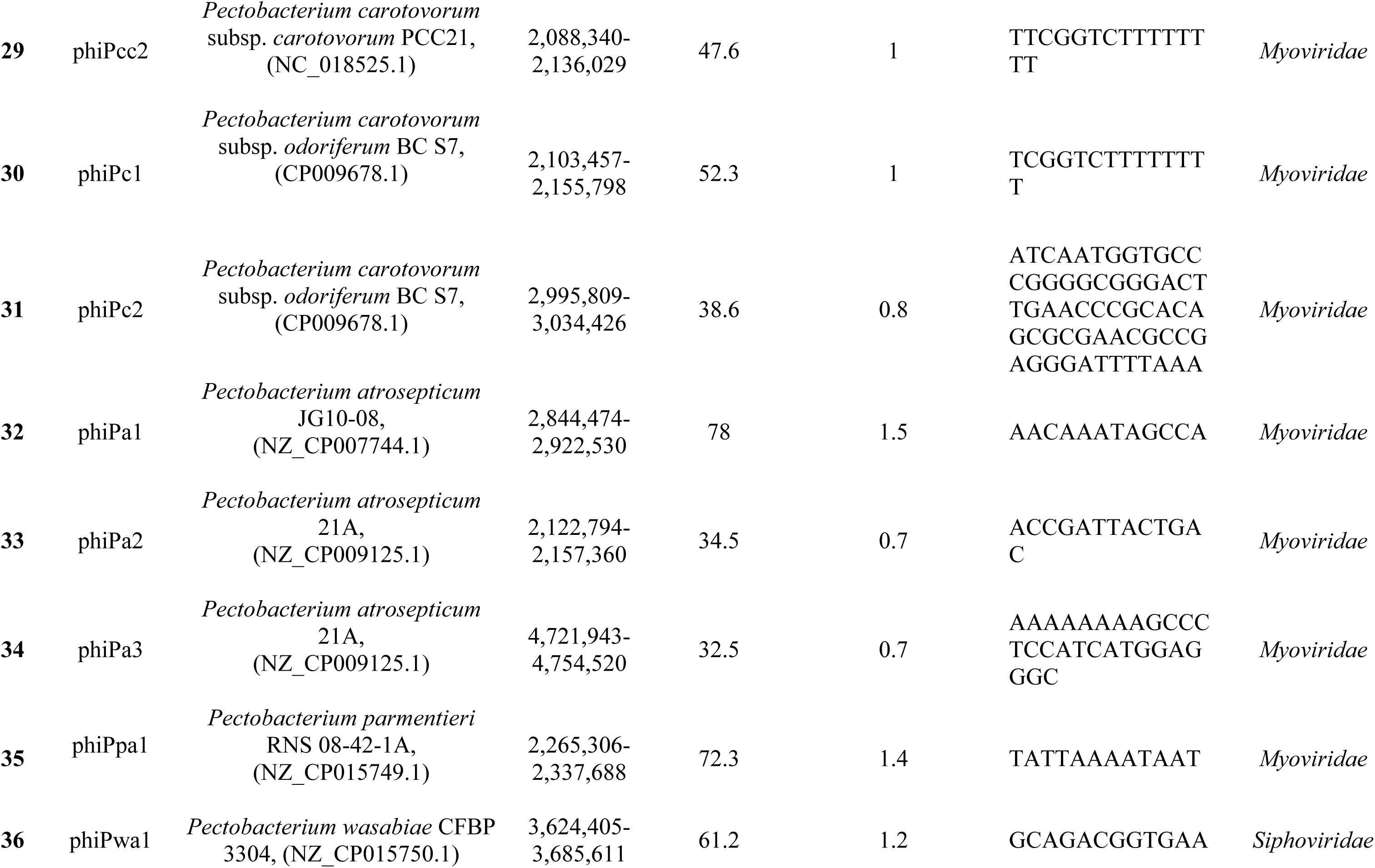
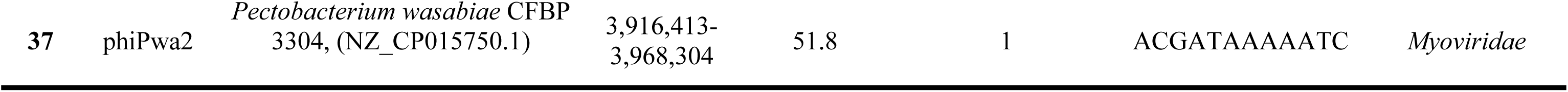
Genomic features of intact prophages present in the complete genomes of Soft Rot *Pectobacteriaceae* (*Dickeya* spp. and *Pectobacterium* spp.) obtained from GenBank (NCBI)

**Figure 1.**
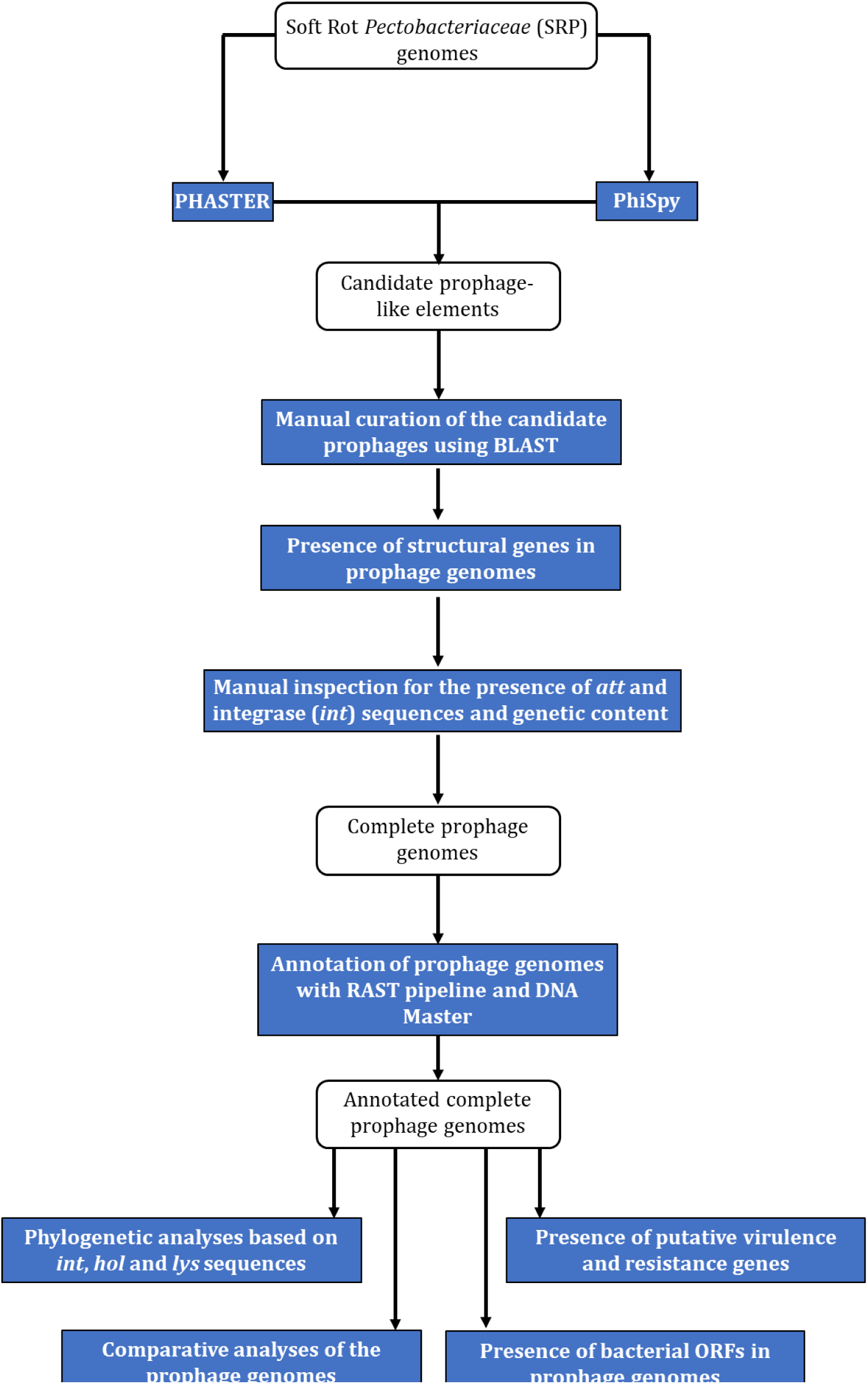
The workflow for identification and characterization of SRP prophages. The blue rectangles represent the tools and methods used for identification of prophages’ sequences in bacterial genomes and the white rounded rectangles represent the data used for the analyses.

### Analyses of prophage genome sequences and comparative genomics

Prophage sequences were annotated using RAST (rast.nmpdr.org) (Aziz et al., 2008;Brettin et al., 2015) (settings: Classic RAST, Glimmer3 release 70, domain Viruses, genetic code:11, disable replication) and DNA Master (Lawrence, University of Pittsburgh, Pennsylvania, USA) (http://en.bio-soft.net/dna/dnamaster.html) using settings advised in (Pope and Jacobs-Sera, 2018). The *attL* and *attR* attachment sites were identified using PHASTER (http://phaster.ca/) (Arndt et al., 2016) and manually inspected using CLC Main Workbench 7 (Qiagen). Multiple sequence alignment of individual prophage genes and phylogenetic analyses were performed using Phylogenetic Pipeline of Information Génomique et Structurale, CNRS-AMU, France (http://www.phylogeny.fr/) and MEGA7 (Kumar et al., 2016). Comparative analyses of the prophage genomes were done using EDGAR (Blom et al., 2009) accessed *via* (https://edgar.computational.bio.uni-giessen.de), DNA Master (Lawrence, University of Pittsburgh, Pennsylvania, USA) (http://en.bio-soft.net/dna/dnamaster.html) and BLASTn accessed *via* (https://blast.ncbi.nlm.nih.gov/Blast.cgi). Pairwise comparison of sequences were analyzed using MAUVE (Darling et al., 2010) and DNA Master. The presence of the putative virulence-associated genes and antibiotic resistance genes in the genomes of prophages was checked using VirulenceFinder ver. 1.5 accessed *via* https://cge.cbs.dtu.dk/services/VirulenceFinder/ and RestFinder ver. 3.0 accessed *via* https://cge.cbs.dtu.dk/services/ResFinder/ (Kleinheinz et al., 2014), respectively.

## RESULTS

### *Presence of prophage-like sequences in* Dickeya *spp. and* Pectobacterium *spp. complete genomes*

The analyses of the fifty seven complete SRP genomes acquired from GenBank (NCBI) with the use of PHASTER and PhiSpy (Fig.1) resulted in discovery of the prophage-like elements in the genomes of 54 SRP strains (in total in 95% genomes) (Fig. 2). Only three *D. solani* genomes, namely *D. solani* strain MK10 (NZ_CM001839.1), *D. solani* strain MK16 (NZ_CM001842.1) and *D. solani* strain GBC 2040 (NZ_CM001860.1) did not harbor any (partial and/or complete) prophage-like elements. Forty eight incomplete (defective) prophage-like elements, i.e. regions that did not contain both attachment sites and sequences coding for integrase were ubiquitously present in the majority of the investigated SRP genomes and varies between 5 kb to 27 kb in size. Often more than one such an element was found per host genome, as it was reported for example in case of *D. chrysanthemi* strain NCPPB 402 (2 defective prophages), *D. dadantii* strain NCPPB 898 (3 defective prophages) and *P. atrosepticum* strain SCRI1043 (2 defective prophages). The subsequent screen of the bacterial genomes for the presence of complete prophage regions yielded 37 complete prophages (27 in *Dickeya* spp. genomes and 10 in *Pectobacterium* spp. genomes) in total in 29 SRP strains (Table 1). Additionally, eight SRP (3 *Pectobacterium* spp. and 5 *Dickeya* spp.) genomes contained more than one complete prophage region (Fig. 2). The sizes of prophage complete genomes varies between 29 kb to 78 kb, an on average these elements comprised between 0.6 to 1.8 % of the host chromosome (Table 1). All the complete prophage genomes possessed structural components related to the phages of the order *Caudovirales* (tailed bacteriophages), which allowed the classification of 30 prophages (81%) to the *Myoviridae* family, 5 prophages (13.5%) to the *Siphoviridae* family and 2 prophages (5.5%) to the *Podoviridae* family (Table 1).

**Figure 2.**
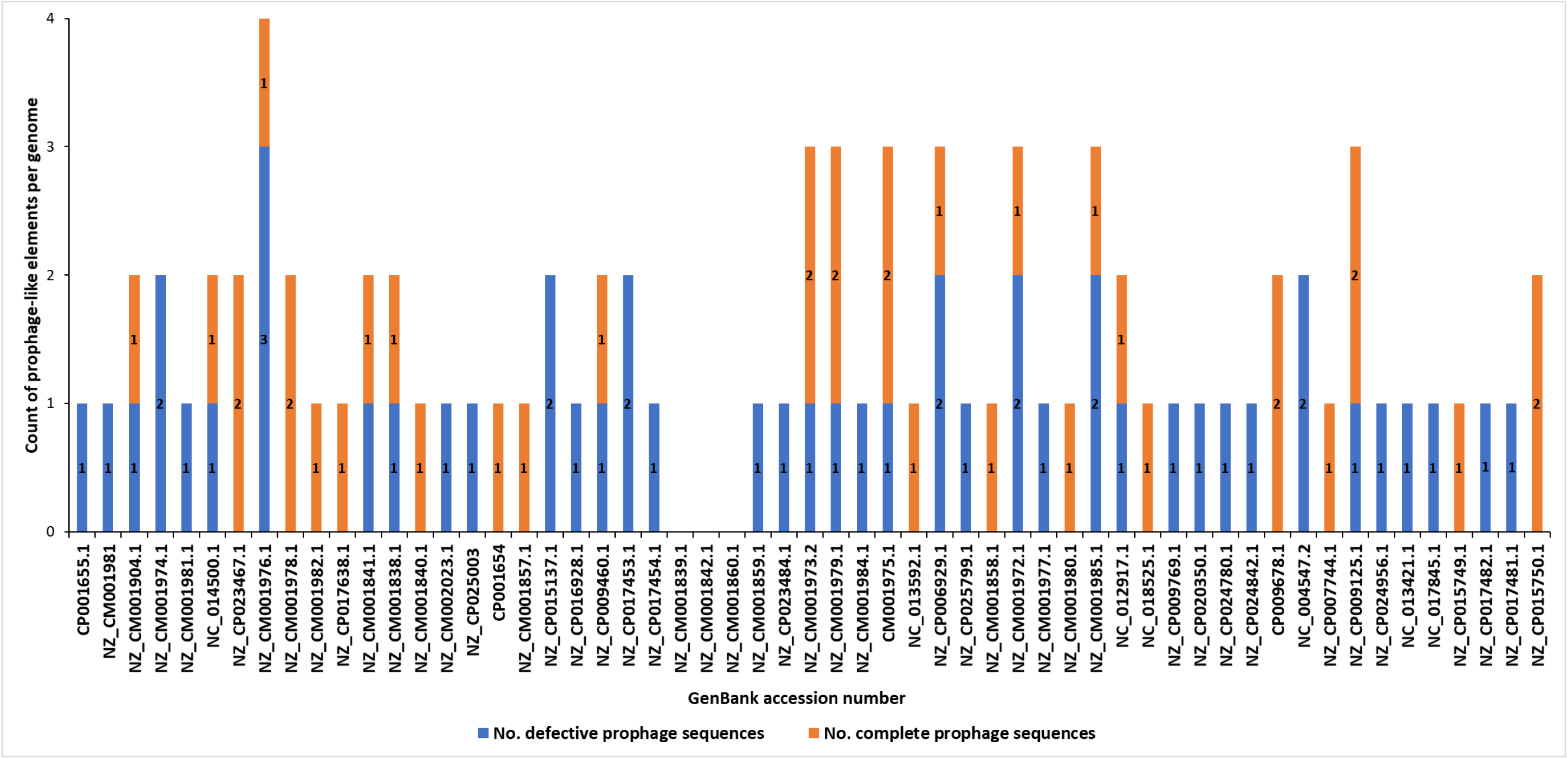
Distribution of defective prophage-like elements and intact prophages in fifty seven complete genome sequences of Soft Rot *Pectobacteriaceae* present in GenBank (NCBI). The prophage sequences were detected using PHASTER and PhiSpy and manually curated using BLAST (NCBI)

### Phylogenetic relationships between prophages found in SRP genomes based on single gene analyses

Integrase, holin and lysin (endolysin, murein hydrolase) amino acid sequences derived from respective nucleotide sequences (*int*, *hol*, *lys*, respectively) found in complete prophage genomes were used to phylogenetically analyze all 37 intact prophages. The *int* gene coding for integrase was present in all 37 screened prophages (Fig. 3A), whereas genes coding for holin (*hol*) and lysin (*lys*) were found inside 21 and 27 prophage sequences, respectively (Fig. 4). The phylogenetic analyses showed that SRP prophages constitute a diverse group; with viruses belonging to the same viral family forming different phylogenetic clades. Likewise, the phylogenetic distance between prophages calculated based on amino acid sequences of integrase had little relevance to the phylogeny between their hosts as no clear separation of the *Pectobacterium* and *Dickeya* prophage clades could be observed (Fig. 3A). In case of the phylogenetic analyses based on amino acid sequences of holin and lysin, prophages present in *Dickeya* spp. genomes were somehow more often constituting clades separated from clades grouping prophages present in the genomes of *Pectobacterium* spp. strains. This separation of clades was however only partial (Fig. 3BC). Based on the integrase and lysin amino acid sequences four prophage clades could be distinguished containing between two and nine prophages each. The use of holin amino acid sequence as a phylogenetic marker allowed differentiation of the three prophage clades (Fig. 3B). Interestingly, the prophages phiDda1, phiDda6, phiD3, phiDdd1, phiDze1 and phiDdi6 were co-grouped both in clade I of holin- and in clade II of integrase-based phylogenetic tries and the prophages phiDze2, phiD5, phiD4 and phiDdd2 were grouped both in clade III of integrase- and clade II of holin-based tries. Likewise, the prophages phiDch1, phiDdi5, phiDdi1 and phiDdi3 were present both in clade III of holin- and clade III of lysin amino acid sequence-based tries.

**Figure 3.**
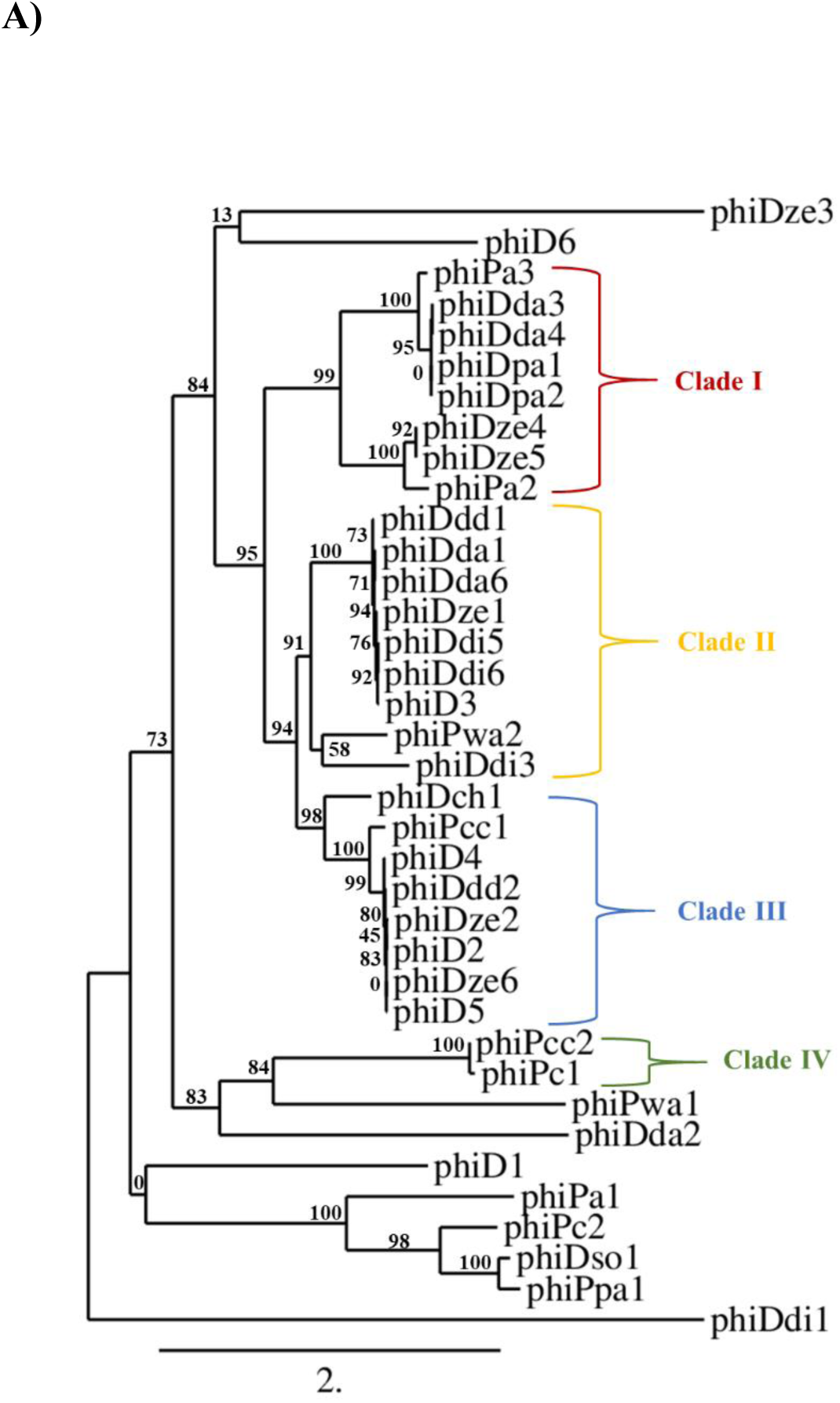
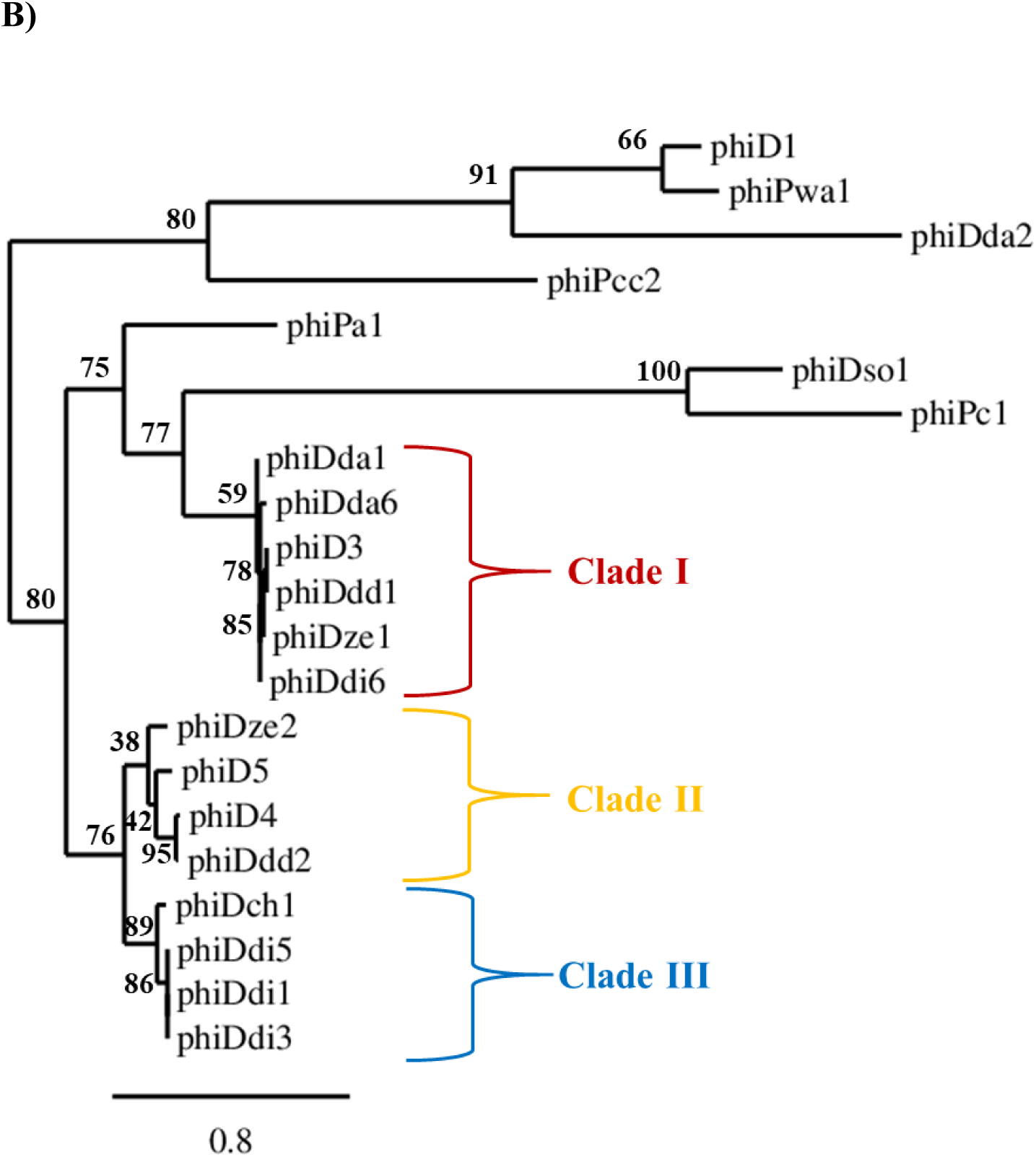
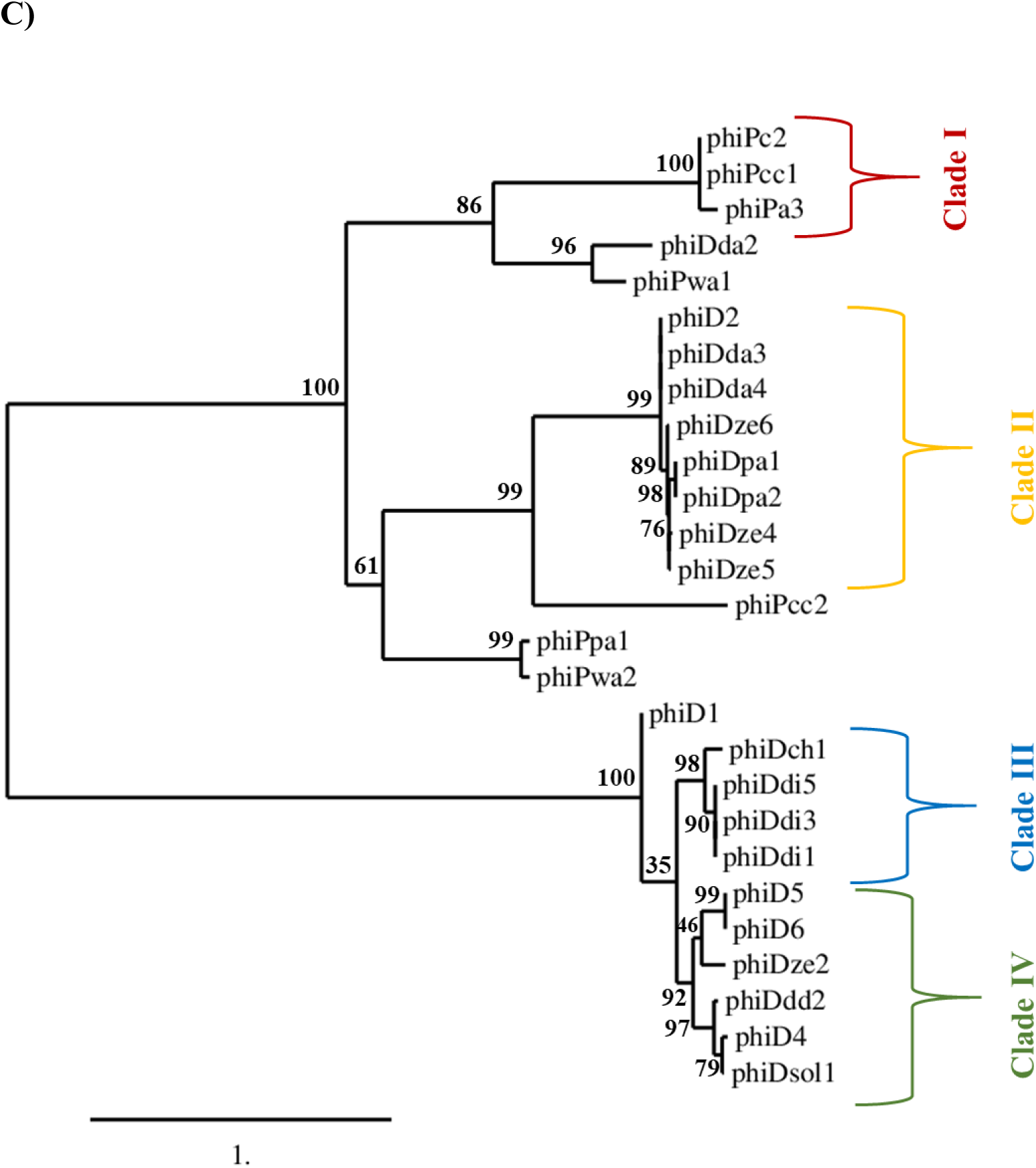
Neighbour-joining (NJ) tree based on the aligned amino acid sequences of integrase (present in 37 prophages) (A), holin (present in 21 prophages) (B) and lysin (present in 27 prophages) (C) genes of intact prophage sequences distributed in 57 Soft Rot *Pectobacteriaceae* genomes. Phylogenetic studies were performed using Phylogenetic Pipeline of Information Génomique et Structurale, CNRS-AMU, France (http://www.phylogeny.fr/) with bootstrap support for 1000 replicates. The bar indicates the number of substitutions per sequence position.

**Figure 4.**
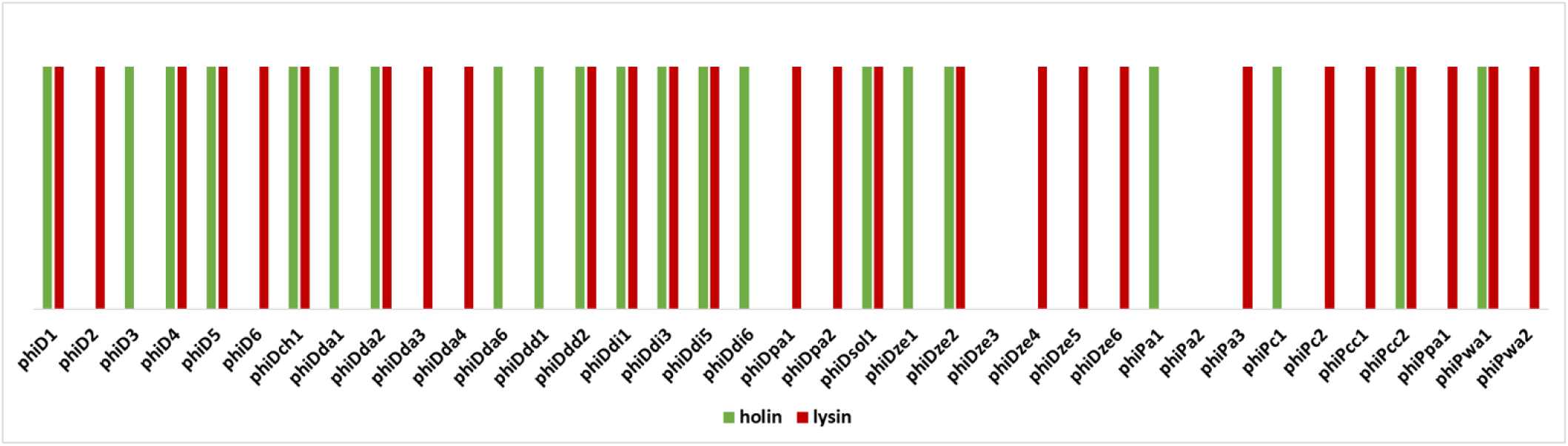
Presence of genes coding for holin and lysin in the genomes of 37 complete prophages. The presence of holin gene is marked with green color and the presence of lysin gene is marked with red color.

### Comparative genomics and proteomics of SRP prophages

Comparative genomics based on the RAST annotated prophage genome sequences allowed visualization of ORFs order present in all 37 prophage genomes (Fig. 5). In general, and with the few exceptions mentioned below, the ORFs organization and arrangement was not conserved over all 37 SRP prophage genomes (high mosaicism). The genome organization was homological only between prophage pairs phiDdi1 and phiDdi3 and highly conserved between prophage pairs phiDze4 and phiDze5 and phiDpa1 and phiDpa2. The phiDdi1 and phiDdi3 were found in two *D. dianthicola* strains namely RNS04.9 and NCPPB 453, prophages phiDze4 and phiDze5 were found in two *D. zeae* isolates CSL RW192 and NCPPB 3531 and prophage phiDpa1 and phiDpa2 were found in two *D. paradisiaca* strains Ech703 and NCPPB 2511. Partial ORFs order conservation was as well present among prophages phiD4, phiD5 and phiDdd2 (Fig 5) residing in *Dickeya* sp. NCPPB 3274, *Dickeya* sp. NCPPB 569 and *D. dadantii* subsp. *diffenbachiae* NCPPB 2976. Frequently, bacterial genome carried two distinct complete prophages as it was identified in the case of *D. dadantii* strain DSM 18020 (carrying prophages phiDda2 and phiDda3), *D. dadantii* subsp. *diffenbachiae* strain NCPPB 2976 (carrying prophages phiDdd1 and phiDdd2), *Dickeya* sp. CSL RW240 (carrying prophages phiD1 and phiD2), *Dickeya* sp. Strain NCPPB 3274 (carrying prophage phiD3 and phiD4), *Dickeya* sp. Strain NCPPB 569 (carrying prophages phiD5 and phiD6) and *P. carotovorum* subsp. *odoriferum* strain BC S7 (carrying prophages phiPc1 and phiPc2). Furthermore, no correlation was observed between host bacterial genome size and prophage genome size (R^2^ = 0.02) (Fig. 6).

**Figure 5.**
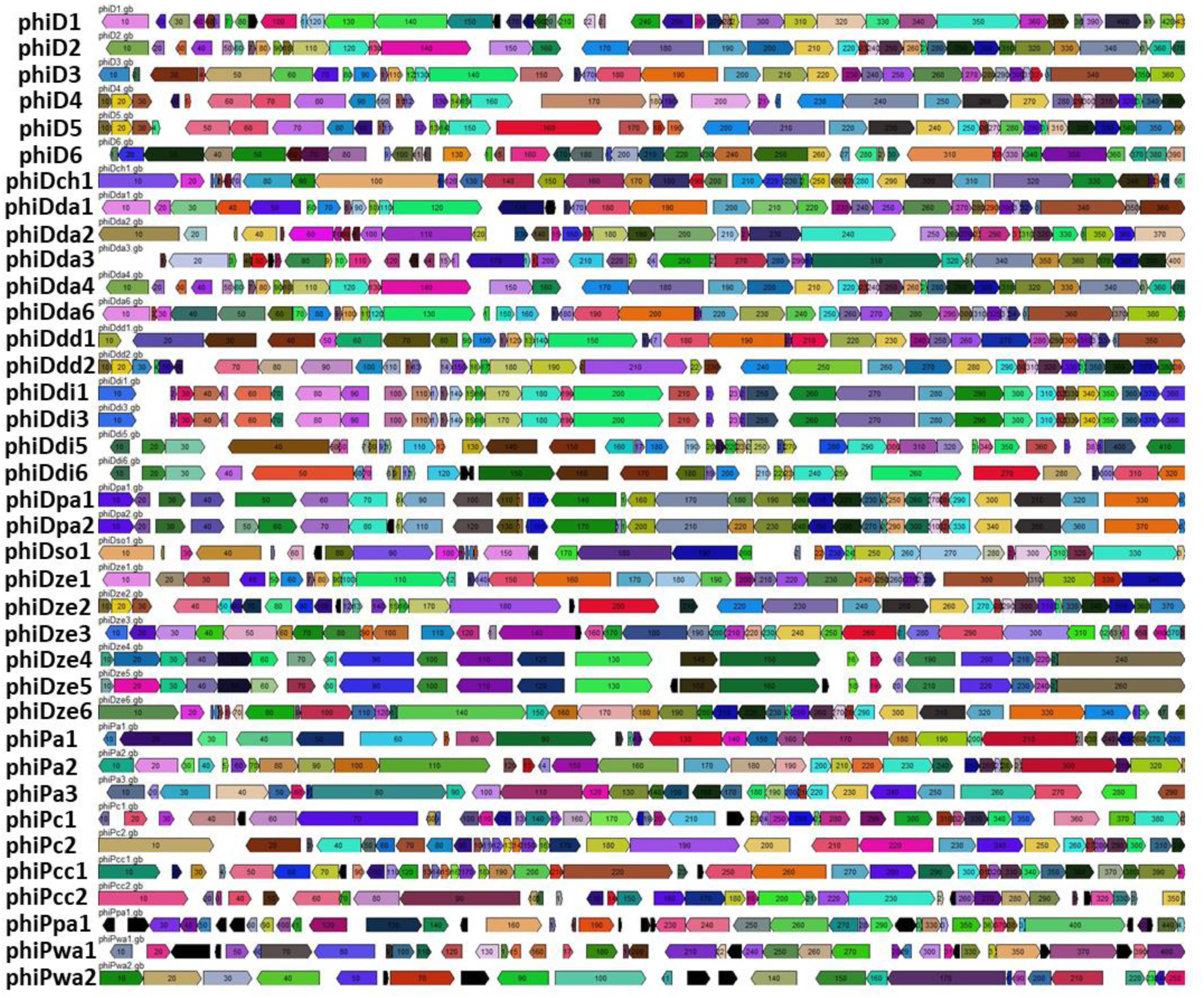
Comparative analyses of the 37 intact prophage genomes. Boxes indicate open reading frames (ORFs) in prophage genomes predicted by RAST annotation pipeline. Homological ORFs are marked with the same color. The analysis and visualization were performed with the use of DNA Master ver. 5.23.2.

**Figure 6.**
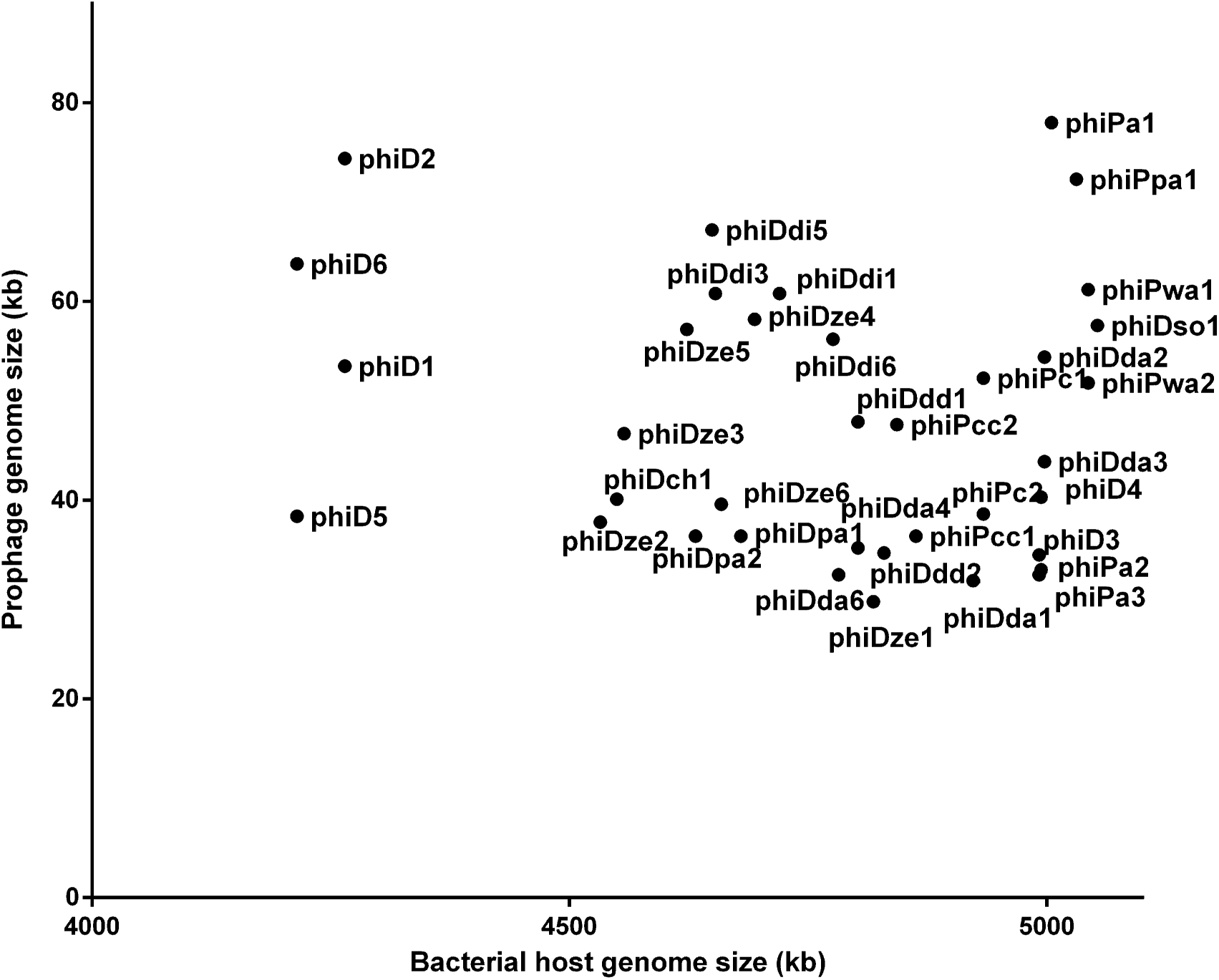
Correlation between the size of the host bacterial genomes and the size of complete prophage genomes. No correlation was found between prophage genome size and host genome size (R^2^ = 0.02).

Dot plot matrix constructed based on average amino acid identity (AAI) of the 37 prophage proteomes resulted in visualization of six distinctive clusters (Fig. 7); two clusters (Cluster 2 and Cluster 3) (Tab. 3 and Tab.4) with the AAI grater than 90%, one cluster with the AAI grater than 85% (Cluster 1) (Tab. 2), two clusters with the AAI grater than 80% (Cluster 4 and Cluster 6) (Tab. 5 and Tab. 7) and one cluster with the AAI grater than 75% (Cluster 5) (Tab. 6) Five clusters (Cluster 1, 2, 3, 4 and 6) were grouping proteomes of prophages present in *Dickeya* spp. genomes, whereas Cluster 5 was grouping prophages hosted by *Pectobacterium* spp. strains.

**Figure 7.**
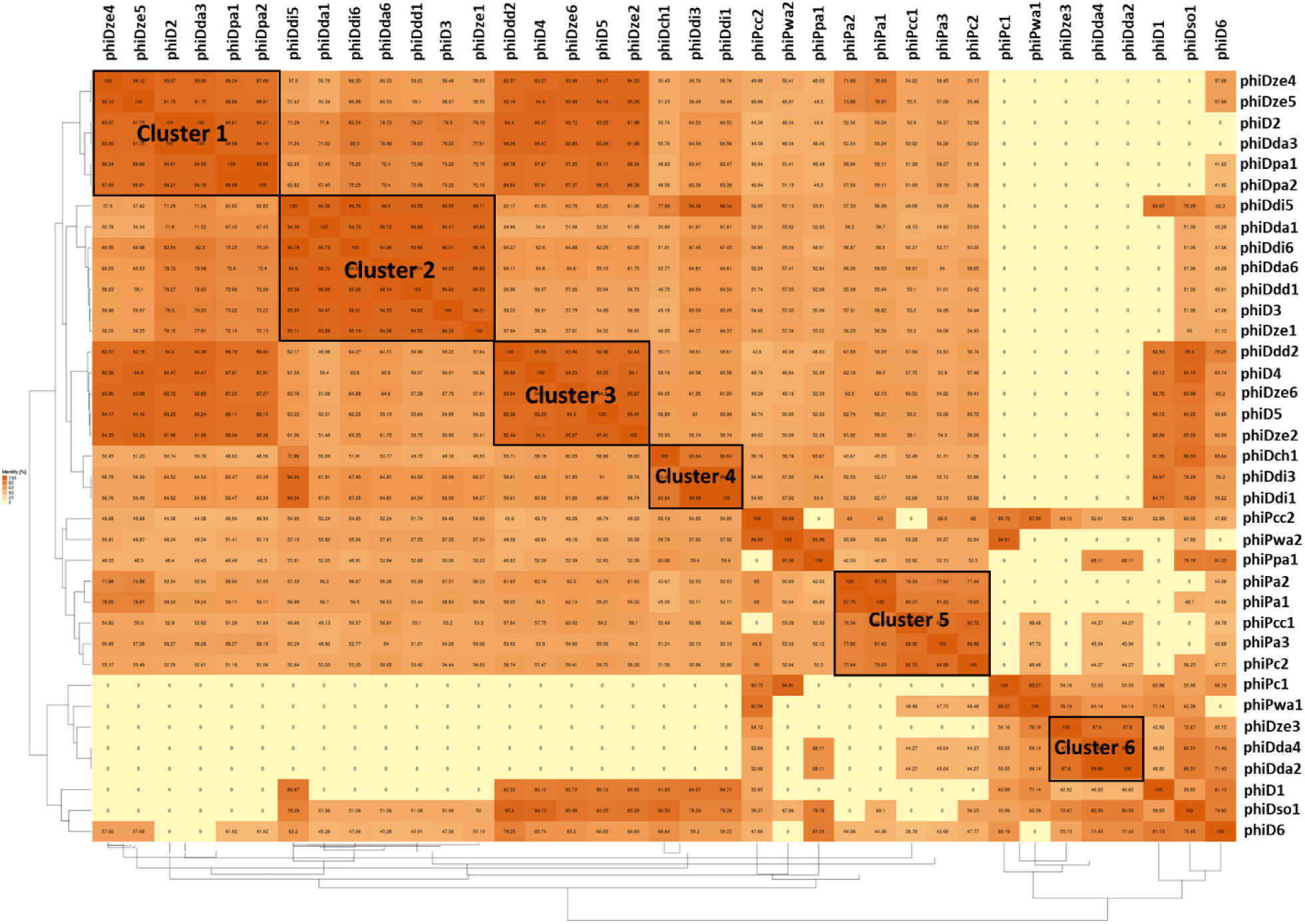
Pairwise average amino acid identity (AAI) heatmap among 37 intact SRP prophages. The map was generated using EDGAR – a 2 software platform for comparative genomics (Blom et al., 2009).

**Table 2.**
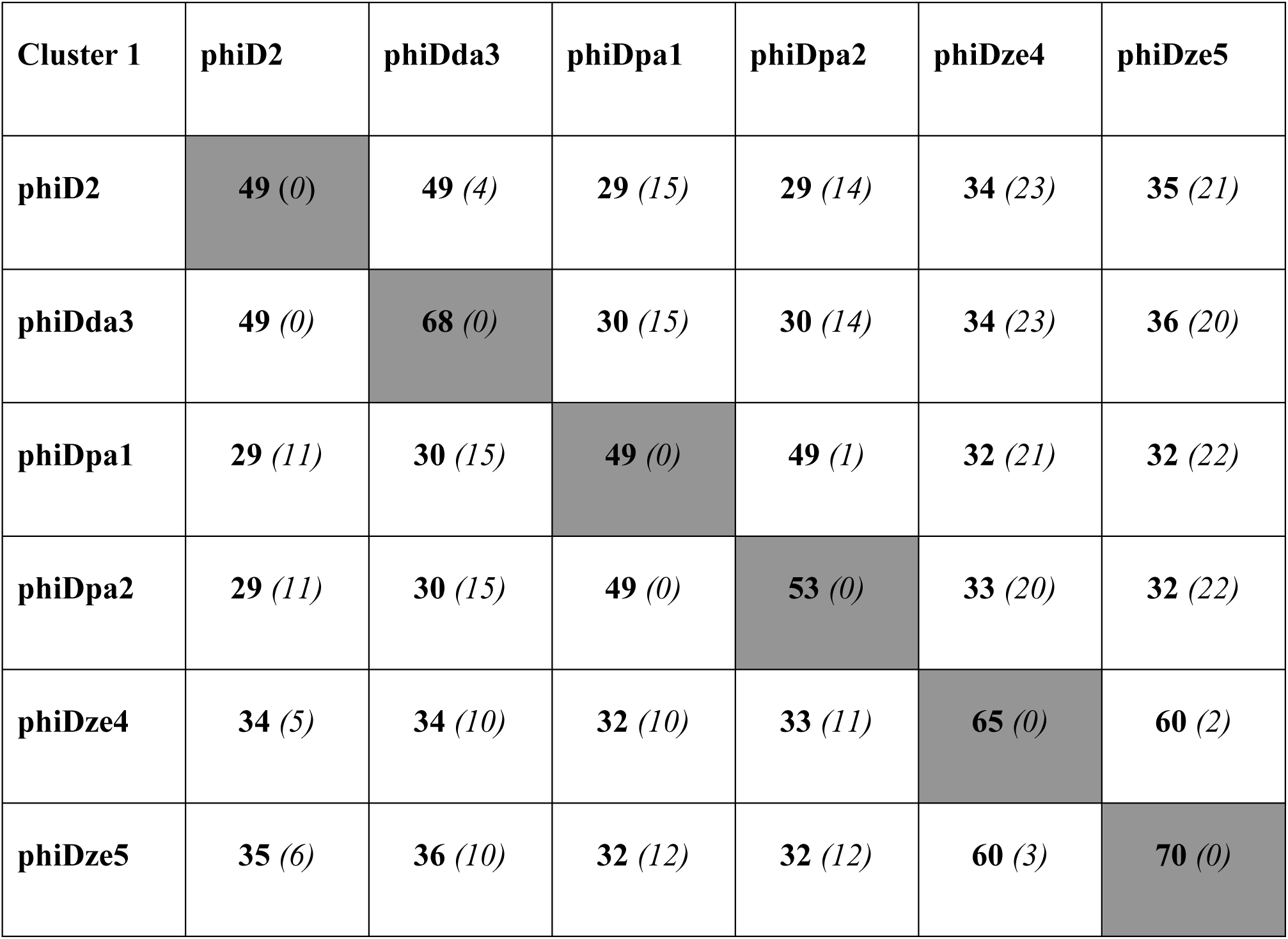
Distinct and shared ORFs present in genomes of prophages: phiD2, phiDda3, phiDpa1, phiDpa2, phiDze4 and phiDze5 constituting AAI Cluster 1. The number of shared ORFs is shown in bold, whereas the number of distinct ORFs is showed in brackets in italic

**Table 3.** Distinct and shared ORFs present in genomes of prophages: phiD3, phiDda1, phiDda6, phiDdd1, phiDdi5, phiDdi6 and phiDze1 constituting AAI Cluster 2. The number of shared ORFs is shown in bold, whereas the number of distinct ORFs is showed in brackets in italic

| Cluster 2 | phiD3 | phiDda1 | phiDda6 | phiDdd1 | phiDdi5 | phiDdi6 | phiDze1 |
| --- | --- | --- | --- | --- | --- | --- | --- |
| phiD3 | **46** (0) | **39** (2) | **42** (4) | **41** (14) | **38** (32) | **39** (16) | **37** (2) |
| phiDda1 | **39** (4) | **44** (0) | **40** (4) | **40** (15) | **36** (34) | **36** (21) | **37** (2) |
| phiDda6 | **42** (3) | **40** (2) | **48** (0) | **42** (14) | **38** (34) | **37** (20) | **37** (2) |
| phiDdd1 | **41** (2) | **40** (1) | **42** (2) | **60** (0) | **50** (23) | **50** (9) | **36** (2) |
| phiDdi5 | **38** (2) | **36** (1) | **38** (2) | **50** (5) | **95** (0) | **66** (3) | **34** (2) |
| phiDdi6 | **39** (1) | **36** (1) | **37** (3) | **50** (5) | **66** (18) | **70** (0) | **35** (2) |
| phiDze1 | **37** (5) | **37** (3) | **37** (5) | **36** (17) | **34** (35) | **35** (20) | **40** (0) |

**Table 4.** Distinct and shared ORFs present in genomes of prophages: phiD4, phiD5, phiDd2, phiDze2 and phiDze6 constituting AAI Cluster 3. The number of shared ORFs is shown in bold, whereas the number of distinct ORFs is showed in brackets in italic

| Cluster 3 | phiD4 | phiD5 | phiDdd2 | phiDze2 | phiDze6 |
| --- | --- | --- | --- | --- | --- |
| phiD4 | **56** (0) | **49** (2) | **42** (2) | **45** (4) | **47** (3) |
| phiD5 | **49** (1) | **55** (0) | **45** (2) | **45** (4) | **49** (3) |
| phiDdd2 | **42** (4) | **45** (4) | **52** (0) | **39** (6) | **43** (5) |
| phiDze2 | **45** (3) | **45** (4) | **39** (4) | **54** (0) | **45** (5) |
| phiDze6 | **47** (1) | **49** (2) | **43** (2) | **45** (4) | **60** (0) |

**Table 5.** Distinct and shared ORFs present in genomes of prophages: phiDch1, phiDdi3 and phiDdi1 constituting AAI Cluster 4. The number of shared ORFs is shown in bold, whereas the number of distinct ORFs is showed in brackets in italic

| Cluster 4 | phiDch1 | phiDdi3 | phiDdi1 |
| --- | --- | --- | --- |
| phiDch1 | **65** (0) | **42** (18) | **42** (18) |
| phiDdi3 | **42** (7) | **81** (0) | **81** (0) |
| phiDdi1 | **42** (7) | **81** (0) | **81** (0) |

**Table 6.** Distinct and shared ORFs present in genomes of prophages: phiPa1, phiPa2, phiPcc1, phiPa3 and phiPc2 constituting AAI Cluster 5. The number of shared ORFs is shown in bold, whereas the number of distinct ORFs is showed in brackets in italic

| **Cluster 5** | **phiPa1** | **phiPa2** | **phiPcc1** | **phiPa3** | **phiPc2** |
| --- | --- | --- | --- | --- | --- |
| **phiPa1** | **81** (*0*) | **39** (*1*) | **22** (*14*) | **24** (*11*) | **24** (*12*) |
| **phiPa2** | **39** (*30*) | **42** (*0*) | **20** (*16*) | **23** (*11*) | **19** (*18*) |
| **phiPcc1** | **22** (*39*) | **20** (*9*) | **52** (*0*) | **32** (*5*) | **39** (*7*) |
| **phiPa3** | **24** (*40*) | **23** (*9*) | **32** (*7*) | **40** (*0*) | **33** (*9*) |
| **phiPc2** | **24** (*37*) | **19** (*10*) | **39** (*4*) | **33** (*4*) | **49** (*0*) |

**Table 7.** Distinct and shared ORFs present in genomes of prophages: phiDze3, phiDda4 and phiDda2 constituting AAI Cluster 6. The number of shared ORFs is shown in bold, whereas the number of distinct ORFs is showed in brackets in italic

| Cluster 6 | phiDze3 | phiDda4 | phiDda2 |
| --- | --- | --- | --- |
| phiDze3 | **82** (0) | **19** (12) | **33** (21) |
| phiDda4 | **19** (34) | **49** (0) | **22** (24) |
| phiDda2 | **33** (26) | **22** (12) | **71** (0) |

### Presence of unique genes of bacterial origin in intact prophage genomes

Of 37 screened complete prophage genomes, only one, phiDze1 did not contain any ORFs of bacterial origin. The other 36 prophages contained between 1 (phiD3, phiDda1) and 23 (phiDdi1 and phiDdi3) ORFs acquired from bacterial hosts (Fig. 8). Most of the bacterial ORFs found in prophages coded for proteins involved in primary bacterial metabolism, proteins associated with DNA/RNA repair, energy transfer, DNA/protein regulation and modification and proteins that may be involved in niche settlement (e.g. coding for resistance to metal ions, nitrogen assimilation, heat shock proteins) (Supplementary Table 1). The most ubiquitously-present among all analyzed intact prophages were: (i) gene coding for methyl-directed repair DNA adenine methylase found in 21 prophage genomes, (ii) gene coding for methyl-transferase found in 9 prophages and (iii) gene coding for modification methylase ScrFIA present in 6 prophage genomes. Interestingly, the similar sets of bacterial genes were found in prophages: phiD2, phiDda3 and phiDda4, prophages phiDdi1 and phiDdi3, prophages phiDdi5 and phiDdi6, prophages phiDpa1 and phiDpa2, prophages phiDze4 and phiDze5 and prophages phiPc2 and phiPcc1 (Supplementary Table 1). Prophages phiD6, phiDdi1, phiDdi1 and phiPa1 carried homological gene encoding tellurite resistance protein TerB, and prophages phiDpa1 and phiDpa2 possesses in their genome gene coding for cation-efflux pump FieF causing resistance to cobalt, zinc and cadmium ions. When screened for putative genes coding for antibiotic synthases, antibiotic resistance genes and genes coding for allergens/toxins, only one prophage, namely phiPpa1 contained gene encoding putative protein involved in biosynthesis of monooxygenase antibiotic. No other such genes were found in the genomes of prophages analyzed by VirulenceFinder and RestFinder and by manual inspection with BLAST.

**Figure 8.**
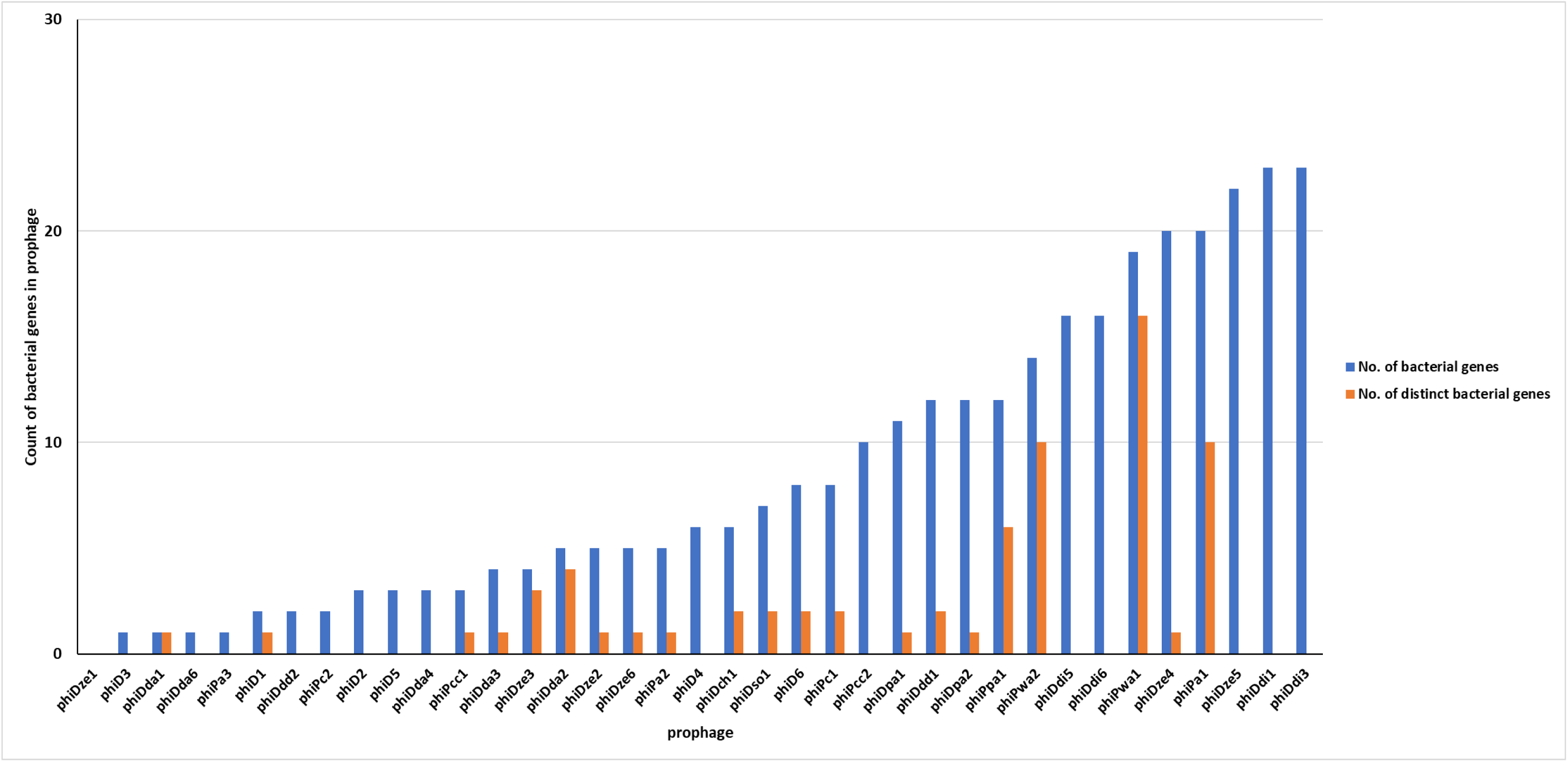
Distribution of genes of bacterial origin in 37 genomes of SRP prophages.

## DISCUSSION

This study was conducted to assess the presence of prophage-like elements in complete genomes of *Pectobacterium* spp. and *Dickeya* spp. strains and to characterize the resulting defective and intact prophages with the use of bioinformatic, comparative techniques. Despite the fact that the majority of bacterial genomes deposited in the international databases contain phage DNA integrated into the host chromosome (Canchaya et al., 2003), the knowledge on prophages inhabiting specifically the SRP genomes is scarce (Varani et al., 2013;Czajkowski, 2015). Likewise, the only temperate bacteriophage described in 1984 and widely used since then for generalized transduction of *Dickeya* spp. is the *D. dadantii* 3937 (at that time named *Erwinia chrysanthemi* 3937) phage phiEC2 analyzed (Resibois et al., 1984). Even this bacteriophage has not been characterized in detail so far and little is known about its ecological, genomic and morphological features (Czajkowski, 2015). The other temperate SPR bacteriophages have been described but to date not characterized on a molecular level (Czajkowski, 2015).

In this study all available fifty seven (August 2018) *Pectobacterium* spp. and *Dickeya* spp. complete genome sequences present in NCBI GenBank were screened for the presence of prophage-like elements. The *in silico* workflow used here (Fig. 1) allowed the identification of prophages in 95% of SRP genomes. Although prophages are known to constitute even as much as 10 to 20% of the bacterium’s genome, the common feature of all prophages analyzed here was that they comprised on average less than 2% of the *Pectobacterium* spp. and *Dickeya* spp. chromosome. It is worth to mention that a food pathogen *Escherichia coli* O157:H7 strain Sakai contains prophages amounted to 16% of its total genome content (Ohnishi et al., 2001). No detailed data exist however on the sizes of prophages present in the genomes of plant pathogenic bacteria and specifically in the genomes of SRP (Varani et al., 2013).

The majority of the SRP prophages (48 sequences) were defective, i.e. did not contain the genes vital for bacteriophage interaction with their hosts such as integrases and genes coding for viral structural proteins. Similarly, some of these sequences were also missing attachment sites. The frequent distribution of incomplete prophages in bacterial chromosomes has been reported for different bacteria including human and animal pathogens as well as for saprophytic bacteria present in soil and water (Casjens, 2003;Bobay et al., 2014). It is widely accepted that bacterial hosts under natural conditions are continuously exposed to phage infections and that some of these events may result in long-term and irreversible phage-bacterial associations on a genomic level (Touchon et al., 2014). This is for sure also true for *Pectobacterium* spp. and *Dickeya* spp. that are characterized by a worldwide distribution (Pérombelon, 2002). The high number of defective prophage sequences reported here may further indicate that the initial rapid inactivation of viable prophages in bacterial genome is followed by a slow deactivation of prophage genes due to the accumulations of point mutants. This so-called phage domestication has been reported for other *Enterobacteriaceae* as a way to cure bacterial genomes from unnecessary and/or toxic genetic material (Bobay et al., 2014).

The 37 complete prophages found in 29 genomes of *Pectobacterium* spp. and *Dickeya* spp. were characterized in detail. Bioinformatic analysis of the prophage genes coding for viral structural proteins allowed classification of these prophages into different families of the order *Caudovirales* (tailed bacteriophages) with the SRP prophages of the *Myoviridae* family being the most abundant (81% of found prophages). The order *Caudovirales* contains more than 97% of all described phages known to infect bacteria with at least 350 distinct phage isolates documented as members of this order to date (Ackermann, 1998;Fokine and Rossmann, 2014). Additionally, more than 99% of all SRP bacteriophages described so far also belong to the order *Caudovirales* and occur in three families namely *Myoviridae*, *Podoviridae* and *Siphoviridae* (Czajkowski, 2015).

The genome organization and ORFs arrangement was not conserved across all 37 intact prophages. The exceptions were the 3 prophage pairs (phiDdi1 and phiDdi3, phiDpa1 and phiiDpa2 and phiDze4 and phiDze5) that were highly conserved with respect to each other. This indicates that overall SRP prophages are possibly mobile, are often transferred between different hosts and easily undergoing rearrangements of their genetic material. It is well established that prophages are highly mosaic and that their genomes consistute modules that can be interchanged between regions and between different phages by recombination (Hendrix et al., 2000). As it is believed now, the constant recombination events and the resulting mosaicism are the major driving force both for bacteriophage and bacterial evolution. On the global scale, it is estimated that for the last 3 billion years, ca. 10^25^ phages initiated infections every second and in each of these incidences bacterial and phage DNA can potentially recombine to generate new genomic arrangements (Hendrix et al., 1999;Pedulla et al., 2003).

No correlation was found between the presence of intact prophages and bacterial genera, bacterial genome size, geographical locations and environments from which the host bacteria were initially isolated. Likewise, due to the absence of universal genes in bacteriophages that can be used for phylogenetic studies (similar to 16S *rDNA* gene in bacteria), (pro)phage classification is difficult (Lawrence et al., 2002). In this study, contrary to the studies performed earlier in which the usefulness of integrase, holin and lysin sequences for the phylogenetic studies of prophages were evaluated (Brüssow et al., 2004;Ventura et al., 2005;Ventura et al., 2007), none of these genes used for phylogenetic analysis of SRP prophages allowed a clear separation of prophages and *Pectobacterium* spp. and *Dickeya* spp. bacteria into distinct clades. Such a phylogenetic diversity suggests an independent evolution of prophages and the SRP bacteria (Colavecchio et al., 2017). This is not of a surprise, taking into account that SRP are not only naturally present in different environments (e.g. soil, water, plant surface, on and inside insects) but are often shifted from one environment to another (Perombelon, 1988;Charkowski, 2006). All these lifestyle changes require a fast adaptation to the new settings (Ma et al., 2007;Reverchon et al., 2016).

More than 50% of complete prophage genomes contained not only the genes coding for structural viral proteins, genes coding for integrases and attachment sites, but also genes coding for holin and lysin. Additionally, the 13 prophages (35% of all complete prophages) contained genes coding for both proteins. Lysins and holins are viral enzymes leading to disruption of the host cells and are facilitating propagation of bacteriophage infection in environment (Wang et al., 2000). As both holin and lysin are essential for host lysis (Young, 2014), the presence of these genes in SRP prophages defines those viruses as infective (Feiner et al., 2015). This further indicates that at least the 13 prophages identified in the genomes of SRP in this work may be easily induced from the host genome and become viable and infectious upon environmental stimuli (Nanda et al., 2014). It may be speculated that induction of SRP prophages will have an impact on environmental fitness and virulence of the hosts. Moreover, the newly found holin and lysin genes, produced at the industrial scale may be a source for the development of bacterial biological control tools for environmental applications as previously suggested (Fenton et al., 2010).

Based on the average amino acid identity (AAI), six prophage clusters; five grouping prophages present in *Dickeya* spp. and one cluster grouping prophages present in *Pectobacterium* spp. could be identified. Opposed to the phylogenetic analyses based on the single prophage gene, AAI seemed to have more power to phylogenetically separate prophages residing in the genomes of *Pectobacterium* spp. and *Dickeya* spp. AAI is suggested to be a better phylogenetic method that can significantly contribute to a whole genome-based taxonomy of *Prokaryotes* (Konstantinidis and Tiedje, 2005), it is however rarely used at the moment to (phylogenetically) analyze viruses.

It is well established that prophages often encode genes (so-called morons) that are not directly involved in viral propagation and infection but that can confer a fitness benefit to their hosts (Bondy-Denomy and Davidson, 2014). Morons can enhance the virulence of the bacteria directly by prophage-encoded toxins and/or indirectly by increasing bacterial fitness which may then results in enhanced virulence (Hacker and Carniel, 2001). All except one of 37 analyzed intact prophages contained genes acquired from host bacteria possible due to the former single and/or multiple infections. Likewise, the most of prophages in this study contained more than one gene of bacterial origin, with two prophages phiDdi1 and phiDdi3 carrying even as many as 23 bacterial genes. Surprisingly, several prophages present in different bacterial genomes carried homological set of bacterial genes indicating that possibly these prophages propagated in co-occurring host populations of different species at the same time. None of the 36 prophages analyzed here however acquired bacterial genes encoding well-described virulence factors exploited by *Pectobacterium* spp. and *Dickeya* spp. to infect plants (Reverchon and Nasser, 2013). The prophages carried genes that may apparently increase hosts ecological fitness in complex and diverse environments; e.g. genes coding for metal ion transporters, enzymes involved in energy metabolism, heat shock proteins, nitrogen assimilation proteins as well as genes coding for DNA methylases which may be used in protecting prophage sequences in the host genome from excision by changing DNA methylation pattern (Canchaya et al., 2003). This may as well explain the high number of prophage sequences observed in many bacterial genomes (Ohnishi et al., 2001;Matos et al., 2013) and relatively higher ratio of prophage-related genes in the pathogenic strains in comparison with saprophytic, non-pathogenic bacteria (Busby et al., 2013). The most commonly present in SRP intact prophage genomes was the gene coding for methyl-directed repair DNA adenine methylase (EC 2.1.1.72) found in 21 viruses. This is a large group of enzymes that apart from being members of restriction-modification systems of many Gram-negative bacteria, play roles in regulation of genes coding for virulence factors in bacterial pathogens at the posttranscriptional level (Marinus and Casadesus, 2009). Unfortunately their roles in pathogenicity of SRPs stands unidentified. It is worth to mention that so far it remains unknown whether the bacterial genes found in this study in 36 prophage genomes undergo transcription and translation.

In general, the relatively high number of intact prophages identified and characterized in this study suggest that the interaction of SRP and bacteriophages in the natural environment may be highly significant for the ecology, adaptation and evolution of *Pectobacterium* spp. and *Dickeya* spp. The prophage induction experiments are now being conducted to further estimate the role of prophages present in SRP strains and to better understand the molecular basis of (pro)phage-bacteria interactions.

## CONCLUSIONS

In the presented study, with the use of bioinformatic tools and manual analyses, for the first time, the 37 intact prophages in the complete genomes of *Pectobacterium* spp. and *Dickeya* spp. were identified and characterized. The prophages, in majority, belonged to the family *Myoviridae* in the order *Caudovirales* and did not possess conserved genome organization (high genetic mosaicism). No correlation was found between the presence of specific prophage and host genome size, bacterial genus and geographical location from which host was isolated. More than half of the analyzed intact prophages were characterized as infectious as they contained genes coding for holin and lysin. Thirty six prophages contained genes of bacterial origin that may increase the ecological fitness of the hosts. To the best of knowledge, these analyses were the first complex comparative studies of the SRP prophages.

## ACKNOWLEDGEMENTS

The work was financially supported by the University of Gdansk, Poland (Uniwersytet Gdanski, Polska) grant no. DS 530-M035-D673-18.

